# Contributions of Low- and High-Frequency Sensorineural Hearing Deficits to Speech Intelligibility in Noise

**DOI:** 10.1101/358127

**Authors:** Sarah Verhulsta, Anna Warzybok

## Abstract

The degree to which supra-threshold hearing deficits affect speech recognition in noise is poorly understood. To clarify the role of hearing sensitivity in different stimulus frequency ranges, and to test the contribution of low- and high-pass speech information to broadband speech recognition, we collected speech reception threshold (SRTs) for low-pass (LP < 1.5 kHz), high-pass (HP > 1.6 kHz) and broadband (BB) speech-in-noise stimuli in 34 listeners. Two noise types with similar long-term spectra were considered: stationary (SSN) and temporally modulated noise (ICRA5-250). Irrespective of the tested listener group (i.e., young normal-hearing, older normal- or impaired-hearing), the BB SRT performance was strongly related to the LP SRT. The encoding of LP speech information was different for SSN and ICRA5-250 noise but similar for HP speech, suggesting a single noise-type invariant coding mechanism for HP speech. Masking release was observed for all filtered conditions and related to the ICRA5-250 SRT. Lastly, the role of hearing sensitivity to the SRT was studied using the speech intelligibility index (SII), which failed to predict the SRTs for the filtered speech conditions and for the older normal-hearing listeners. This suggests that supra-threshold hearing deficits are important contributors to the SRT of older normal-hearing listeners.

## 1. Introduction

**Abbreviations**
PTA: pure tone average
SRT: speech reception threshold
AM DT: amplitude-modulation detection threshold
BB: broadband
LP: low-pass
HP: high-pass
LF: low-frequency
HF: high-frequency
NH: normal-hearing
HI: hearing-impaired
SSN: speech-shaped noise
ICRA: modulated International Collegium of Rehabilitative Audiology noise
TFS: temporal fine-structure
TENV: temporal envelope

Older normal-hearing and hearing-impaired listeners often experience difficulties in understanding speech when it is presented in background noise. When seeking help for these problems, audiological practice mostly assesses hearing status using the pure-tone audiogram or speech recognition in quiet. There are several reports showing that these standard clinical measures are poor predictors of speech recognition in noise (Festen and Plomp 1983; Papakonstantinou, Strelcyk, and Dau 2011), and that supra-threshold hearing deficits targeting temporal fine-structure (TFS) or envelope encoding (TENV) are important for speech intelligibility as well (Lorenzi et al. 2006; Hopkins and Moore 2011; Papakonstantinou, Strelcyk, and Dau 2011; Léger, Moore, and Lorenzi 2012a,b). Lastly, there is growing physiological evidence that supra-threshold processing of audible sound might be compromised by synaptopathy (Bharadwaj et al. 2014, 2015; Liberman et al. 2016; Parthasarathy and Kujawa 2018; Parthasarathy, Herrmann, and Bartlett 2018) as a consequence of aging.

As it is not clear what the respective contribution of hearing sensitivity or supra-threshold deficits to speech recognition is, we predicted individual SRTs for each listener by means of the speech intelligibility index (SII; ANSI 1997) which only considers individual audibility deficits. Failing to account for individual differences on the basis of the SII alone, would indicate there are other deficits contributing to speech-in-noise recognition as well. Secondly, it is crucial to understand how different frequency regions and associated hearing deficits *contribute* to broadband speech intelligibility to develop effective hearing restoration strategies. Peripheral hearing damage is frequency-specific and may entail degradation in different coding mechanisms (e.g., audibility, TFS and TENV encoding).

Our study design compared SRTs of low-pass (LP) and high-pass (HP) filtered stimuli with SRTs of broadband speech-in-noise stimuli to quantify these contributions. The cut-off frequency of the LP and HP filter separates the speech material in equal band-importance functions (1.5 kHz; ANSI 1997) and was further motivated by our understanding of the physiology of hearing which roughly associates LP speech recognition with TFS coding ability (Léger, Moore, and Lorenzi 2012b; Nelson et al. 2003; Füllgrabe, Berthommier, and Lorenzi 2006; Hopkins and Moore 2009), and HP speech recognition with robust envelope encoding in the presence of background noise (Carney, Li, and McDonough 2015; Henry, Kale, and Heinz 2016). The filtered stimuli were not amplified, in contrast to traditional studies who amplified HP filtered speech to render the speech material equally audible in normal and hearing-impaired test groups, or to maintain an equal loudness percept for LP & HP filtered stimuli (e.g. Lorenzi et al. 2009; Hopkins, Moore, and Stone 2008; Goossens et al. 2017). The amplification approach is appropriate for investigating whether supra-threshold hearing deficits persist when audibility is restored, but they are not able to clarify which frequency regions and associated hearing deficits are responsible for degrading the SRT to broadband speech in normal listening conditions. Our study design (which did not amplify the LP or HP filtered broadband speech and noise material at the tested SNRs) can characterize whether LP and HP frequency regions equally contribute to the SRT to broadband speech, while quantifying to which degree hearing sensitivity in LP or HP frequency bands can account for the measured SRT.

Lastly, we investigated how encoding of LP and HP speech is affected by the temporal characteristics of the noise envelope. The temporal characteristics of modulated noise are known to result in lower (better) SRTs than for stationary noise (Miller and Licklider 1950; Takahashi and Bacon 1992; Festen and Plomp 1990; Gnansia, Jourdes, and Lorenzi 2008). This *masking release* is reduced or even absent in older normal-hearing (Dubno, Horwitz, and Ahlstrom 2002; Grose, Mamo, and Hall III 2009; Goossens et al. 2017) or hearing-impaired listeners (Wagener 2003; Versfeld and Dreschler 2002), but it is not clear which mechanisms drive this degradation or whether LP and HP frequency regions contribute equally to broadband masking release. Taken together, this study aims at better understanding the contribution of hearing sensitivity and supra-threshold hearing deficits to speech recognition in stationary or modulated noise and characterizes the contribution of speech information in LP and HP frequency regions to broadband stimuli in stationary and modulated noise. We focus on normal-hearing younger and older listeners as well as mildly-impaired older listeners to cover a range of participants with age or hearing-sensitivity related hearing damage. The study outcomes are important for the development of effective hearing loss compensation strategies.

## 2. Methods

### 2.1. Study Participants

Fourteen young normal-hearing (yNH, 19-29 yr), 10 older normal-hearing (oNH, 56-77 yr), and 10 older hearing-impaired listeners (oHI, 61-78 yr) were selected to participate in the study. Participants received written and oral information about the experiments, gave written consent, and were paid for their participation. Experiments were approved by the ethics committee of Oldenburg University. The two older listener groups were age-matched (p>0.05) with a mean age of 68.4±7.2 years. The yNH group had a mean age of 24.6±3.1 and differed significantly in age from both oNH and oHI groups (p<0.001). Mean pure-tone audiograms with corresponding standard deviations are shown for each listener group in Fig.1A. A repeated-measures ANOVA with audiometric frequency as a within-subject factor and listener group as a between-subject factor yielded a significant effect of frequency [F(3.15, 97.8)=83.37, p<0.001 Greenhouse-Geisser correction)], listener group [F(2, 166)=116, p<0.001] and interaction [F(6.31, 97.8)=17.2, p<0.001]. Post-hoc comparisons for each tested frequency (Bonferroni corrected) did not show significant differences in pure-tone detection threshold between yNH and oNH listeners for frequencies up to 2 kHz (p>0.05, Fig.1A), whereas significant group differences were found for higher frequencies (p<0.05). The hearing thresholds of the oNH listeners did not exceed 20 dB HL in the frequency range that is most important for speech perception (0.125-4 kHz). Pure-tone thresholds significantly differed for all frequencies between yNH and oHI groups (p<0.05), and for frequencies > 500 Hz between oNH and oHI groups. The pure-tone average over 0.5, 1, 2, and 4 kHz (PTA_0.5,1,2,4_), averaged across listeners was 3.3±1.3, 7.6±2.8, and 25.8±6.4 dB HL for the yNH, oNH, and oHI groups, respectively. Figure 1B shows the relationship between PTA_0.5,1,2,4_ and age.

**Figure 1.**
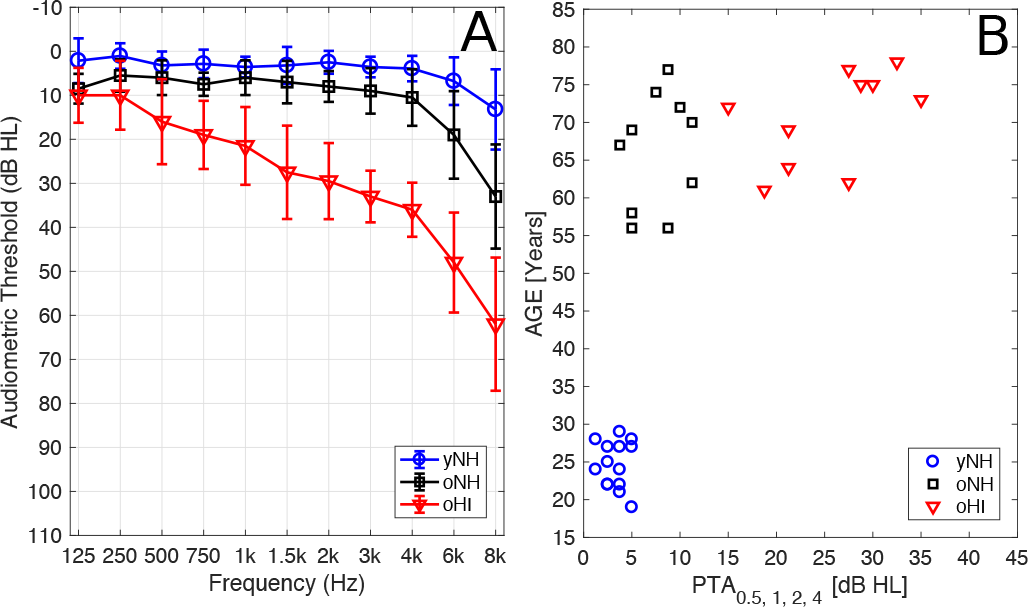
**A**: Mean pure tone thresholds with corresponding standard deviations for groups of young normal-hearing (NH, blue, n=14), older normal-hearing (oNH, black, n=10), and older hearing-impaired listeners (oHI, red, n=10). **B**: Relation between the participants’ age and pure tone average (PTA0.5,1,2,4).

### 2.2. Speech Recognition

A 1-up/1-down adaptive procedure with varying step size and word scoring was used to determine the SRT corresponding to 50% correctly identified words. The Oldenburg sentence test (Wagener, Kühnel, and Kollmeier 1999; Brand and Kollmeier 2002b) with test lists of 20 sentences was used in each condition. Speech recognition measurements included: *(i)* SRT in quiet, *(ii)* SRT in speech-shaped noise (SSN) with broadband (BB), low-pass (LP), and high-pass (HP) filtered stimuli, and *(iii)*, SRT in modulated noise (ICRA5-250; Wagener, Brand, and Kollmeier 2006) using BB, LP, and HP filtered stimuli. Both speech and noise were filtered in the LP and HP conditions. The SSN was generated by multiple superpositions of the test material so that the long-term spectrum of the SSN matched that of the sentences. The ICRA5-250 noise matched the envelope of a single male talker and includes pause durations up to 250 ms. The long-term spectrum of the ICRA5-250 noise was shaped according to the international long-term average speech spectrum (Wagener, Brand, and Kollmeier 2006). Previous studies showed that the long-term spectrum of the SSN derived from the Oldenburg sentence test and the spectrum of the ICRA1 noise (stationary noise with the same long-term spectrum as ICRA5-250) resulted in the same SRTs (Hochmuth et al. 2015; Wagener and Brand 2005). We thus assume that any differences in SRT observed between SSN and ICRA5-250 cannot be explained on the basis of spectral differences between the two maskers. Prior to the measurements, all listeners were trained with two test lists of BB stimuli and one test list with the HP filtered stimuli, all presented in SSN.

The LP and HP condition were generated by filtering the speech and noise signals using a 1024^*th*^ order FFT filter with cut-off frequencies of 1.5 and 1.65 kHz, respectively. The cut-off frequencies of the filters were based on dividing the speech signals in two parts of comparable importance, considering the SII band importance function for speech in noise (ANSI 1997; Hochmuth et al. 2015). The sum of the SII octave band importance weights is 0.457 for frequencies 0.25, 0.5, and 1 kHz (roughly corresponding to the LP condition in this study) and 0.543 for frequencies 2, 4, and 8 kHz (roughly corresponding to the HP condition in this study). In the BB SSN and ICRA5-250 conditions, the noise level was fixed at 70 dB SPL while the speech level was varied adaptively to determine the SRT. For all conditions tested, the filtered stimuli were not amplified compared to the BB condition to ensure that the BB and filtered stimuli had the same spectral level in the pass band. This method allows for the determination of the contribution of specific frequency regions to BB speech recognition, but resulted in lower overall levels of the filtered speech and noise. The difference in overall level was ≈ 0 dB between the BB and LP conditions, and ≈19 dB between the BB and HP conditions. Previous studies have shown that for the Oldenburg sentence test the SRT is not influenced by the absolute level of background noise (Wagener 2003). Therefore, it is assumed that the absolute level difference between BB and HP condition should not affect SRTs. Measurements were conducted in a double-walled sound-insulated booth and the sentences were presented over free-field equalized Sennheiser HDA200 headphones. The speech recognition experiments were administered using the Oldenburg Measurement Platform (HörTech gGmbH) with the Earbox Highpower ear 3.0 sound card. The measurement setup was calibrated using Brüel & Kjær instruments (artificial ear type 4153 connected with microphone type 4134, preamplifier type 2669, and amplifier type 2610).

### 2.3. Pearson Correlations, Regression Models and Multiple Comparisons

We considered Pearsons’ *r* correlations for metrics collected from all listeners in the study (pooled across yNH, oNH and oHI; n=34), as well as for subgroups of NH (yNH+oNH; n=24) and older listeners (oNH+oHI; n=20). *r*-values and their respective p-values were only reported when the variables in the correlation passed the Kolmogorov-Smirnov test for normality. p-values were Bonferroni corrected on the basis of all speech and MR conditions (total of 10), resulting in significant correlations when p<0.005. Post-hoc t-tests which investigated the effect of listener group of a specific variable were Bonferroni corrected by the number of test groups (i.e. p<0.016).

In a few occasions throughout the study, it was necessary to compare *r* obtained for one variable *z* and two independent variables (*x* and *y*) to draw conclusions about which variable related more strongly to the individual results in the *z* variable. A significance value which explains whether the *r* value between *x* & *z* was significantly stronger than the *r* between *y* & *z*, was computed numerically using a resampling technique (Wilcox 2009). N *x-z* and *y-z* data-point pairs were drawn (with replacement) from the total N pairs and served to calculate *r*_x-z,n_ and *r*_y-z,n_. 200 of such draws were taken for each *x-z* and *y-z* dataset to obtain two normal distributions of *r*_x-z_ and *r*_y-z_ values. These were entered in a standard t-test to yield a p-value.

While the participant groups were selected to yield three homogeneous groups with similarly shaped audiograms or ages, it is also interesting to study whether individual differences in performance can be explained by individual differences in PTA or age to isolate near-threshold from supra-threshold deficits in degraded speech recognition. Multiple regression models with PTA and age as explaining variables were applied to study this aspect, but they could only be applied for the older listener group (oNH+oHI, n=20) for which PTA and age were normally distributed. Multiple regression models for the pooled group of all listeners or the normal-hearing (yNH+oNH) subgroup were not meaningful as either age or PTA was not normally distributed in these subgroups.

### 2.4. Speech Intelligibility Index Simulations

The SII was computed by dividing the long-term speech and noise spectra into different frequency bands and estimating the weighted SNR average across frequency bands. We used the implementation of Beutelmann, Brand, and Kollmeier (2010), for which the number and bandwidths of the SII filter bands were adapted according to the auditory gammatone filter bank (Hohmann 2002). The basic calculation procedure of the SII followed the description in the ANSI standard (ANSI 1997). To account for hearing impairment in the SII calculation, an internal noise was derived from the individual audiograms and added to the external noise. For yNH listeners, the use of individual audiograms does not significantly influence the accuracy of speech intelligibility predictions in noise and the observed variance in measured SRTs can thus not be predicted by the model (Beutelmann, Brand, and Kollmeier 2010). An average audiogram of 0 dB HL was hence assumed for the yNH listeners. Individual audiograms were fit to the group of oNH individuals with mild high-frequency losses and the oHI individuals. The SRT for a given condition was calculated by selecting a fixed reference SII value and by varying the SNR until the SII matched the reference. The BB stationary noise condition with SRT of yNH listeners was defined as the reference condition. The speech signal consisted of 10 concatenated OLSA sentences and the extended version of the model was applied (short-time SII; Beutelmann, Brand, and Kollmeier 2010) as this model was proposed for SII predictions with modulated or time variant interferers. The short-time SII analyses the signals in short time frames of 1024 samples and adopts a 44100 Hz sampling rate with a half frame-length overlap. The effective frame length was 12 ms and our parameters corresponded to those adopted in previous studies (Rhebergen, Versfeld, and Dreschler 2006; Beutelmann, Brand, and Kollmeier 2010). To predict SRTs for the filtered conditions, two SII approaches were tested. The first used the original SII with all frequency bands, the second used only frequency bands which were represented in the filtered signals.

## 3. Results

Mean SRT and masking release values as well as their standard deviations are summarized in Table1 for the yNH, oNH and oHI listener groups.

### 3.1. Speech Recognition in Quiet

Statistically significant differences were found between the listener groups [F(2,33)=44.5, p<0.001]. Post-hoc comparisons with Bonferroni correction confirmed the significant SRT differences across groups. Fig.2 shows that hearing sensitivity, as characterized by PTA_0.5,1,2,4_ was strongly related to the SRT and age when speech was presented in the absence of noise and all listeners were included in the analysis. These results are in line with previous studies (e.g. Festen and Plomp 1983; Smoorenburg 1992). The SRTs of the yNH group corroborated earlier reports (Brand and Kollmeier 2002a; Wagener 2003; Meister et al. 2011) The SRTs for the oNH group were on average 6.4 dB higher than for the yNH. It is not clear whether age-related or PTA deficits were responsible for the degraded oNH performance. However, since oHI listeners performed worse than oNH listeners (of the same age), we suggest that PTA, and not age-related, differences were responsible for the group differences between yNH, oNH and oHI speech recognition in quiet.

**Figure 2.**
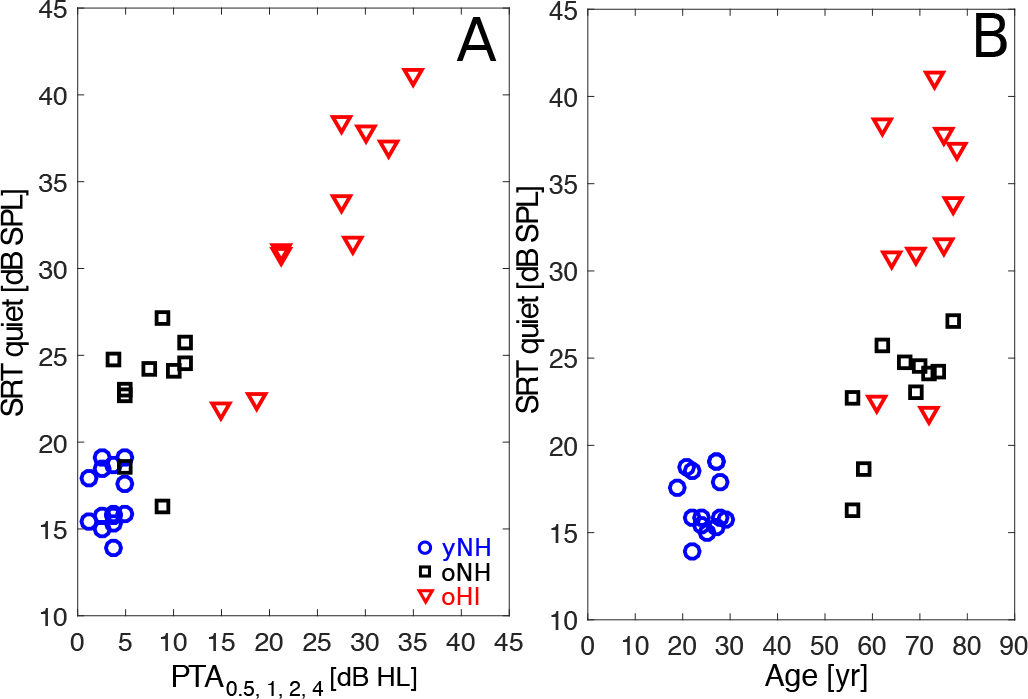
Relation between the individual SRT in quiet and PTA0.5,1,2,4 (**A**) or age (**B**).

Further, within the subset of older listeners (oNH+oHI), for whom both age and PTA_0.5,1,2,4_ were normally distributed, PTA_0.5,1,2,4_ differences explained the individual SRTs. The linear regression between PTA_0.5,1,2,4_ and SRT was significant (d.f.=18, p=3.15e-7), and after accounting for age in a multiple regression model (d.f.=17, 1.38e-6), PTA_0.5,1,2,4_ remained the dominant factor (p=4.8e-6) over age (p=0.2).

### 3.2. Speech Recognition in Noise

There were significant between-group SRT differences in the SSN_BB_ condition [F(2,33)=6.1, p=0.006]. A post-hoc t-test with Bonferroni correction for test group, showed that the yNH group (SRT=−6.4 dB) scored significantly better than oNH (SRT=−5.4 dB, p=0.034) and oHI (SRT=−5.3 dB, p=0.01) groups (Fig.3A). Within the subgroup of age-matched older listeners, neither age, nor PTA_0.5,1,2,4_, nor a combination of both, showed significant relationships to the SRT in linear regression models, suggesting another origin for the individual differences. As both oNH and oHI listeners performed significantly worse than yNH listeners and no significant differences were found between the oNH and oHI group performance, it appears that a general age-related effect degraded the SSN_BB_ SRT of the older subjects. Consequently, which individual differences within age-matched older listeners appeared no longer linearly related to age or PTA differences.

**Figure 3.**
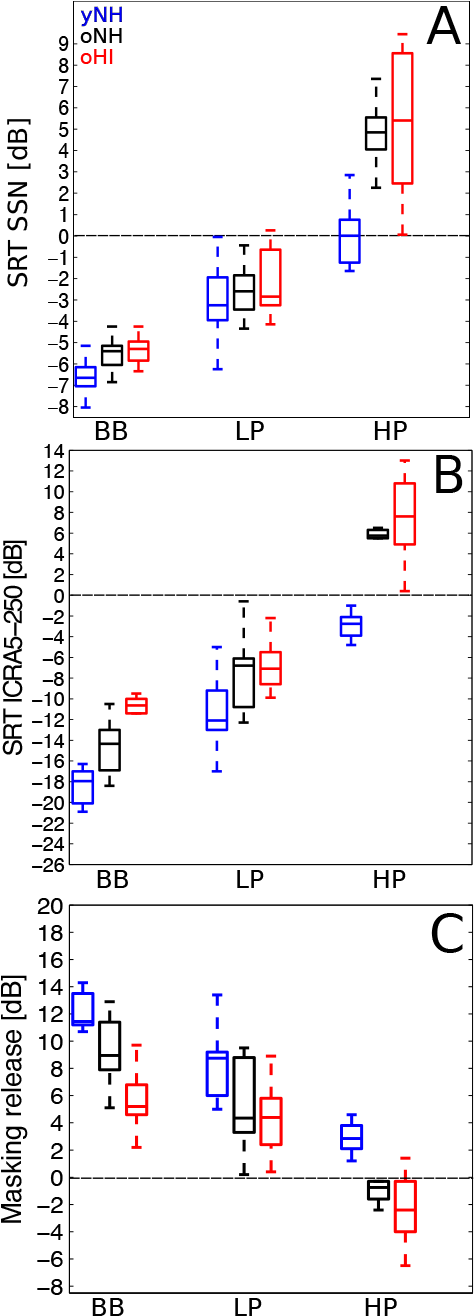
Boxplots of SRTs in the three participant groups (yNH, oNH, oHI) for broadband (BB), low-pass (LP) and high-pass (BB) filtered speech conditions. **A**: SRT in SSN, **B**: SRT in ICRA5-250 noise, **C**: Masking release defined as the SRT in ICRA5-125 noise - SRT in SSN.

SRTs in modulated noise (ICRA5-250) differed across groups (Fig.3B; [F(2,33)=29.4, p<0.001]) and post-hoc comparisons (Bonferroni corrected for group) revealed significant differences across all three groups (p<0.01). The ICRA_BB_ SRT for yNH listeners was −18.8 dB and degraded to −14.6 dB and −10.9 dB for oNH and oHI listeners, respectively. In contrast to the SSN_BB_ condition, individual ICRA_BB_ SRTs within the older group (oNH+oHI) related both to age (p=0.035) and PTA_0.5,1,2,4_ (p<0.001) in single regression models, and to PTA_0.5,1,2,4_ (p=0.006) when age was accounted for in a multiple regression model. Hearing sensitivity thus plays a bigger role in explaining individual performance among older listeners in the ICRA_BB_ than the SSN_BB_ condition.

### 3.3. Masking Release Benefit

ICRA5-250BB SRTs were on average 9.5 dB better than SSNBB SRTs. This SRT difference is called masking release (MR), and was significantly different across groups (F(2,31)=156, p=0.001; Fig.3C). The oNH and oHI groups benefited on average 3.2 and 6.7 dB less from the modulated noise than the yNH group. The lower amounts of MR for oNH listeners are in line with several other studies which reported reduced MR in older NH listeners (Dubno, Horwitz, and Ahlstrom 2002; Grose, Mamo, and Hall III 2009; Goossens et al. 2017). Lastly, within the age-matched older group (oNH+oHI), reduced amounts of MR were strongly related to reduced hearing sensitivity (r_older_, BB=0.73, p<0.001), corroborating earlier reports (Festen and Plomp 1990; Strelcyk and Dau 2009; Christiansen and Dau 2012). The linear regression between MRBB and PTA0.5,1,2,4 (d.f.=18, p=0.036) and MRBB and age were significant (d.f.=18, p=2.5e-4), but after accounting for age in a multiple regression model (d.f.=17, 9.6e-4), PTA0.5,1,2,4 remained the dominant factor (p=0.002) explaining individual differences over age (p=0.33). In summary, a general age effect was seen to reduce MR in the older listener groups, after which PTA0.5,1,2,4 differences were responsible for explaining the individual performance within an age-matched older group.

### 3.4. Role of Audibility in SRTs and Masking Release

Fig.4 shows scatter plots of measured and predicted SRTs in SSN, ICRA5-250 noise, as well as masking release. A strong correlation between measured and predicted data indicates that individual differences across listeners are well predicted by individual hearing threshold differences. In contrast, a weak correlation indicates that the differences in SRT across listeners cannot be explained on the basis of hearing threshold differences and that other deficits influence performance. Audibility accounted for 57% of the ICRA_BB_ SRT variance (with an RMS error between measured and predicted SRTs of 3.6 dB) but failed to explain individual differences in SSN_BB_ SRT (r_all_=0.09). MR_BB_ differences were well captured by the SII model (r_all_=0.62, RMS error of 2.3 dB). Considering the NH (yNH+oNH) group, no correlation was found between measured and predicted data. This indicates that performance differences between the yNH and oNH group cannot be explained on the basis of pure threshold differences.

**Figure 4.**
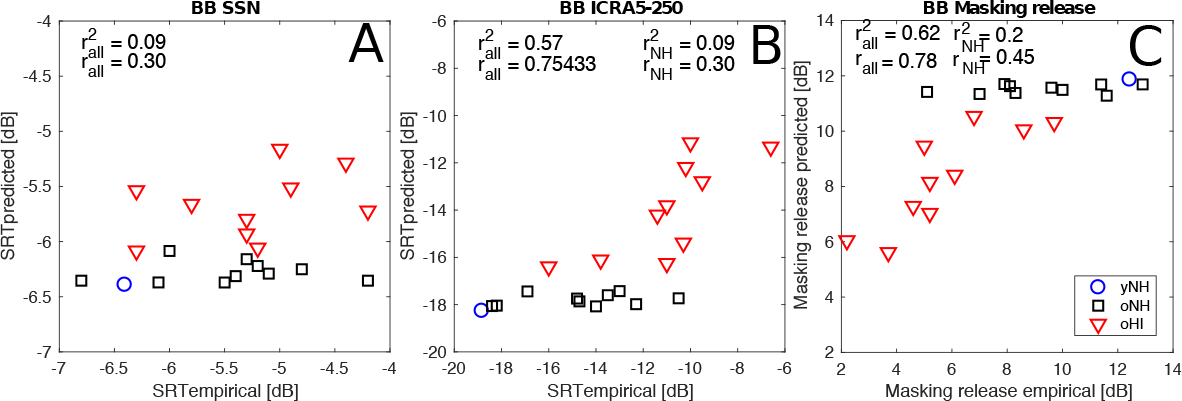
Scatter plot of empirical and predicted broadband SRTs in the stationary (SSN, left) and modulated (ICRA5-250, middle) noise as well as broadband masking release (right) for yNH (circles), oNH (squares), and oHI (downward-pointing triangles) listeners.

### 3.5. Contribution of LP and HP SRTs to the SSN_BB_ SRT

By comparing SRTs in the SSN_BB_ condition to LP and HP filtered SSN conditions, we attempted to assess which frequency regions and associated hearing deficits determined performance in the BB condition. Figure 3 and Table 1 summarize the mean results. For the SSN condition: *(i)* HP SRTs showed an effect of listener group [F(2,33)=18.0, p<0.005; Fig.3A] that was absent for the LP SRTs [F(2,33=1.3), p>0.05]. *(ii)* The average LP SRT was 3 dB worse than the BB SRT, while a larger SRT drop of 9 dB was observed for the HP condition (Fig.3A). *(iii)* A correlation and resampling analysis between the filtered and BB conditions (Fig.5) indicated that the BB SRT was more strongly correlated to the LP SRT (r_all_= 0.77; Fig.5A) than to the HP SRT (r_all_=0.61; Fig.5B), suggesting a stronger contribution of LP cues to the BB SRT (resampling analysis comparing r’s: p<0.001). Lastly, *(iv)*, individual differences in the HP condition were not predictive (r=0.36, p>0.05) of performance in the LP condition for NH listeners (Fig.5C), indicating there are different speech-in-SSN coding mechanism within the LP and HP frequency regions in the absence of PTA deficits.

**Table 1.** Mean SRTs and MRs with standard deviations for the tested conditions and listener groups: yNH (n=14), oNH (n=10) and oHI (n=10).

|  | quiet<sub>BB</sub> | SSN<sub>BB</sub> | ICRA<sub>BB</sub> | MR<sub>BB</sub> |  | SSN<sub>LP</sub> | ICRA<sub>LP</sub> | MR<sub>LP</sub> |  | SSN<sub>HP</sub> | ICRA<sub>HP</sub> | MR<sub>HP</sub> |
| --- | --- | --- | --- | --- | --- | --- | --- | --- | --- | --- | --- | --- |
| yNH | 16.7±1.7 | -6.4±1.0 | -18.8±2.4 | 12.4±2.4 |  | -3.15±1.7 | -11.35±3.2 | -8.2±3.3 |  | 0.3±1.8 | -2.7±1.8 | 2.9±2.3 |
| oNH | 23.1±3.3 | 9.2±2.6 | -14.6±2.6 | -5.4±0.7 |  | -2.53±1.2 | -7.7±3.5 | 5.17±3.5 |  | 4.7±1.5 | 5.48±1.5 | -0.74±2.9 |
| oHI | 32.6±6.5 | -5.3±0.7 | -10.9±2.5 | 5.7±2.5 |  | -2.2±1.5 | -6.6±2.5 | 4.4±2.5 |  | 5.4±3.4 | 7.6±3.4 | -2.2±2.4 |

**Figure 5.**
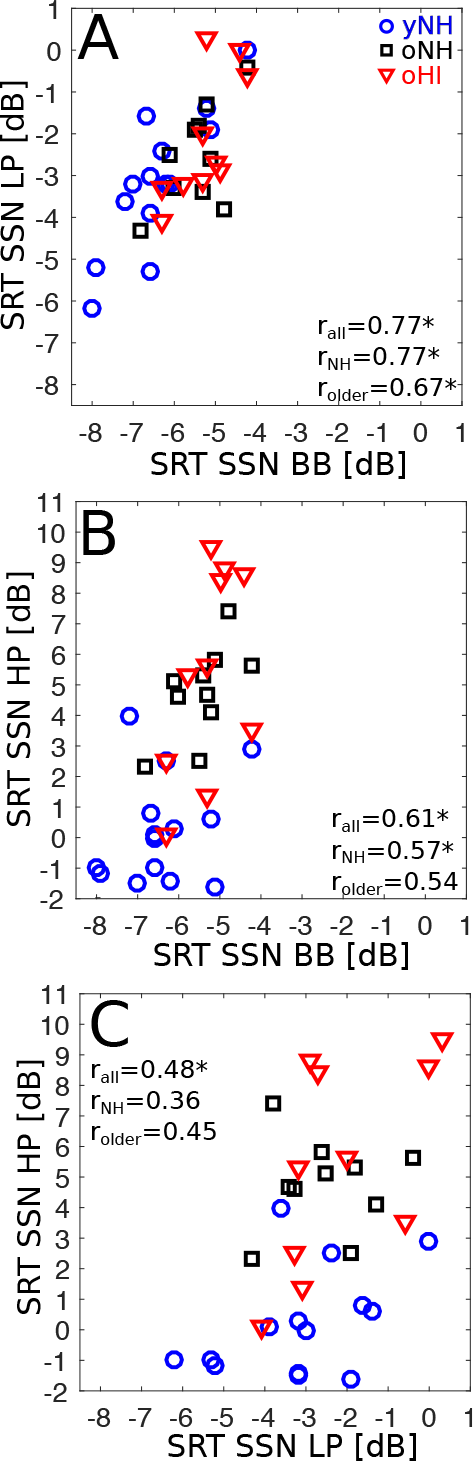
Relations between SRTs of the BB SSN noise condition and the filtered LP and HP conditions. *r* is given for all participants together (*r*_all_), and for subgroups of normal-hearing (*r*_NH_; yNH+oNH), and older listeners (*r*_older_; oNH+oHI). Significance is indicated by a* for the Bonferroni corrected of p=0.005.

oNH listeners performed significantly worse in SSN_HP_ (SRT=4.7 dB) than the yNH group (SRT=0.25 dB, p<0.005) and similarly to the oHI group (SRT=5.4 dB, p>0.99), even though the latter group had elevated PTA_2,4_s. Within the older group (oNH+oHI), neither age nor PTA threshold significantly affected the SSN SRT. Neither single or multiple linear regression models for SSN_LP_ or SSN_HP_ with age and/or PTA as the explaining variable reached significance. Factors different from PTA and age where thus responsible for individual performance differences for older listeners with normal or impaired audiograms.

### 3.6. Contribution of LP and HP SRTs to the ICRA5-250_BB_ SRT

Compared to the ICRA5-250_BB_ SRT of −18.8 dB, the LP and HP filtered conditions yielded 6.2 dB and 17.7 dB higher SRTs (resulting in positive SNRs), respectively. This implies that the LP region made an overall stronger contribution to the ICRA5-250_BB_ SRT. However, Fig.6 shows that the HP SRT was more strongly correlated to the BB SRT (r_all_, HP=0.85; Fig.6B) than the LP SRT (r_all_, LP=0.70, Fig.6A) and the resampling analysis confirmed that the r values were significantly different p<0.001). This suggests that even though the average LP SRT was closer to the average BB SRT, high-frequency cues dominated the individual SRT variation for the ICRA_BB_ condition. This observation is opposite to the trend observed for the SSN condition, for which the LP SRT explained the variance of the BB SRT better than the HP SRT. Lastly, the relationship between the LP and HP filtered ICRA5-250 SRTs was investigated in Fig.6C. Neither the correlations in the NH (yNH+oNH) or older (oNH+oHI) subgroups reached significance at p<0.005, suggesting the existence of different speech-in-noise coding mechanisms in the two frequency regions.

**Figure 6.**
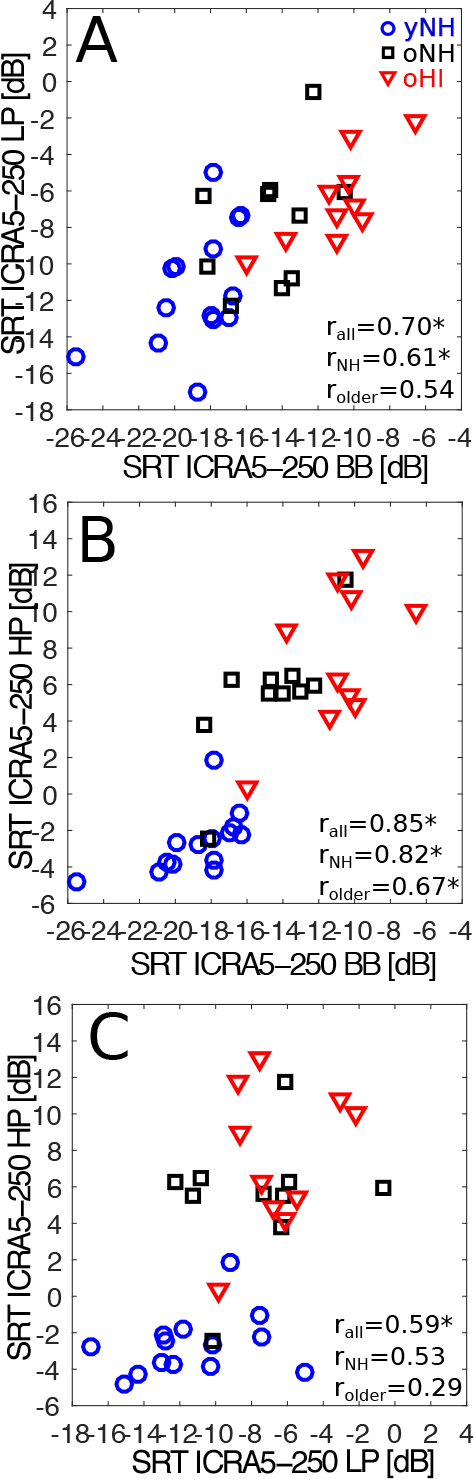
Relations between SRTs of the BB ICRA5-250 noise condition and the filtered LP and HP conditions. *r* is given for all participants together (*r*_all_), and for subgroups of normal-hearing (*r*_NH_; yNH+oNH), and older listeners (*r*_older_; oNH+oHI). Significance is indicated by a * for the Bonferroni corrected of p=0.005.

yNH listeners performed consistently better than older listeners in both LP and HP filtered ICRA conditions (p<0.001) and no significant differences were found between the oNH and oHI group (p>0.05) who significantly differed in hearing sensitivity, but not in age (Fig.3B, table 1). ICRA_LP_ SRTs were neither related to PTA_0.5,1_ nor age within the older group (oNH+oHI). Both single and multiple regression models with these variables failed to reach significance. Differently, the ICRA_HP_ SRTs were both predicted by PTA_2,4_ (d.f.=18, p=0.027) and age (d.f.=18, p=0.003). After accounting for PTA_2,4_ (p=0.31) in a multiple regression model, only age (p=0.0032) remained as an explaining variable (d.f.=17, p=0.001). This suggests that age-related factors (and not hearing sensitivity) play a an important role in determining the ICRA_HP_ SRT in an age-matched older group with normal or impaired audiograms.

### 3.7. Contribution of LP and HP masking release to MR_BB_

Band limiting the stimuli in the LP and HP conditions significantly affected MR (F(2,31)=156, p<0.001; Fig.3C and Table 1). The drop in MR was 4.2 dB for the LP condition and 9.4 dB for the HP condition compared to the BB MR of 12.4 dB (yNH group). Most of the older listeners gave negative MR values in the HP condition, which showed a strong relationship to age (r_oNH+oHI_=−0.71, p<0.001). The added modulation can thus be detrimental for HP speech recognition in older listeners and result in a form of *modulation interference*. Modulation interference was particularly pronounced for the MR_HP_ condition. A multiple regression model for age and PTA_2,4_ confirmed that increased modulation interference related to the listeners’ age within the older listener group, after accounting for hearing sensitivity (model p=0.002). This relationship was not observed for MR_LP_, for which neither age or PTA_0.5,1_ explained individual differences in the (positive) masking release values of the older listeners.

Lastly, we investigated whether the MR_BB_ equalled the sum of the MRs obtained for the low and high-frequency regions. MR_BB_ related strongly to the sum of MR_LP_ and MR_HP_ (r_all_=0.72, p<0.001) suggesting that both frequency regions contribute to providing MR to speech in broadband noise. The additivity of MR across frequency bands also implies that negative MR_HP_s impair the MR_BB_, even if there is a MR_LP_ benefit.

### 3.8. Mechanisms Involved in LP and HP Speech Encoding

To investigate the role of audibility in the LP and HP conditions, we first calculated the SII for the filtered conditions and compared the results to the SRTs of the yNH listeners. Figure 7 shows that the SII predictions failed to account for the empirical SRTs: the original SII implementation predicted similar SRTs in all SSN and ICRA conditions (BB, LP, HP). The modified model captured the increasing trend of SRTs from the BB to LP and HP conditions better, but predicted larger SRTs for the LP than for both masker types. Given that the SII models failed to predict the baseline NH SRT in the filtered conditions, we did not report results for oNH or oHI listeners.

**Figure 7.**
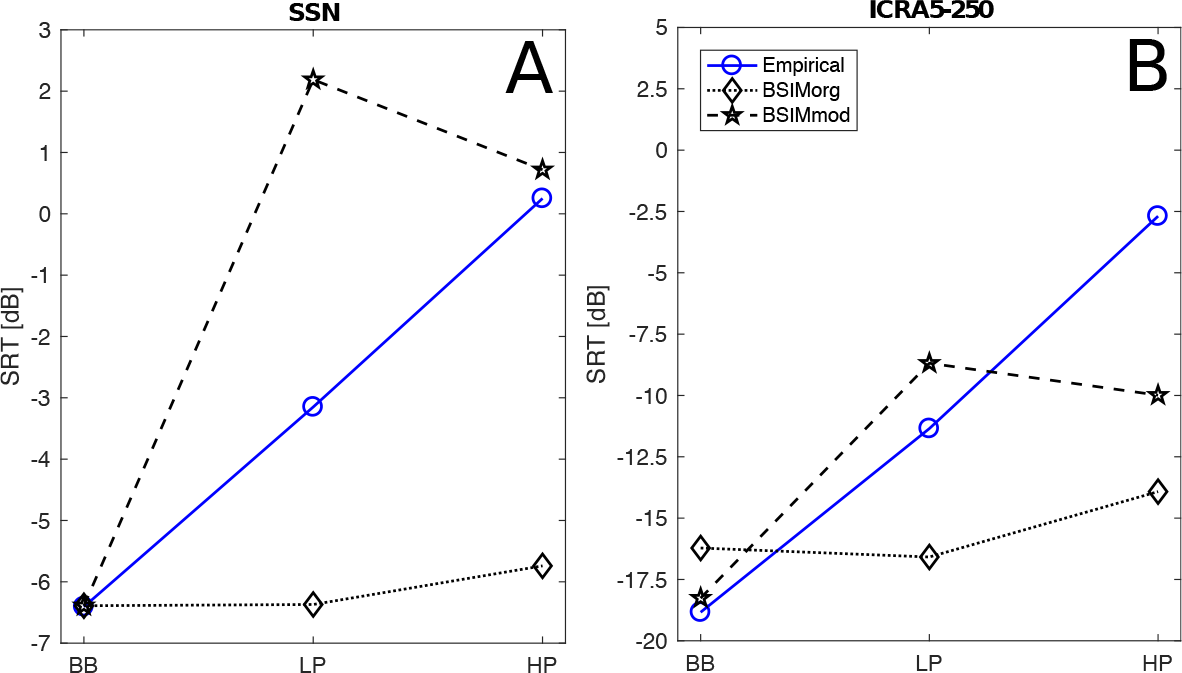
Measured (circles) and predicted SRTs in the stationary (SSN, left) and modulated (ICRA5-250, right) noise for broadband (BB), low-pass (LP), and high-pass (HP) filtered stimuli and young normal-hearing listeners. Diamonds and pentagrams indicate predictions with the original and modified SII implementation, respectively.

In the absence of SII predictions for the filtered conditions, we plotted SSN and ICRA SRTs against each other in Fig.8 to further explore which hearing mechanisms are important at the considered frequency regions. Within the NH group, SSN_LP_ and ICRA_LP_ SRTs were related (r_NH_ = 0.65, p<0.005), whereas this relationship was absent among participants in the older group (r_old_ = 0.41, p>0.05). Hearing deficits in this age-matched older group did not equally affect performance in modulated or SSN noise, whereas this appeared the case for the younger group who had age, but not threshold, differences.

The SSN_HP_ and ICRA_HP_ SRTs (Fig.8B) related well to each other (r_all_ = 0.83, p<0.005), both in the NH and older listener group (r_NH_ = 0.79, p<0.005 and r_older_ = 0.71, p<0.005, respectively), suggesting that HF hearing deficits (irrespective of their origin) affected speech recognition in modulated and stationary noise similarly.

### 3.9. Discussion

This study investigated how LP and HP portions of speech-in-noise stimuli contribute to BB speech-in-noise recognition and showed a dominant role for LP (<1.5 kHz) over HP speech recognition, irrespective of the considered group or noise type (Fig.3). This is somewhat surprising given that the LP and HP cut-off frequencies were chosen to maintain a similar weight in the SII band-importance function for speech (ANSI 1997). This discrepancy might relate to the construction of the band-importance function, which emphasizes the high-frequency content of sentence material (i.e., maximum weight on the 1-2 kHz region). Because it was recently shown that SII predictions for the German Matrix test in noise were significantly worse than the empirical data for listeners with steeply sloping audiograms (rms error of 22.6 dB) (Hülsmeier et al. 2018), there may be a general overestimation of high-frequency content in the SII.

#### 3.9.1 Individual Characterization of Speech Recognition Deficits

Our study suggests that LP mechanisms are important predictors of individual BB speech recognition in a fixed 70-dB-SPL SSN (Fig.5A), and this has implications for the use of speech-in-noise tests for hearing diagnostics and hearing-loss treatment: *(i)* Using SSN_BB_ SRTs to diagnose hearing ability informs mostly about low-frequency hearing deficits. *(ii)* Hearing restoration strategies aimed to enhance high-frequency (>2 kHz) speech cuese in older or hearing-impaired listeners may not improve the SSN_BB_ SRT, as even for a full restoration (or in the absence of hearing deficits) the SSN_HP_ SRT is about 3 dB higher than the SSN_LP_ SRT in yNH listeners (Fig.3A).

The noise type influenced which frequency regions and associated hearing deficits the speech-in-noise test was most sensitive to. Individual ICRA5-250_HP_ SRTs were a better predictor of ICRA5-250_BB_ SRTs than were ICRA5-250_LP_ SRTs (Fig.6), suggesting that the ICRA5-250_BB_ SRT test may be more sensitive to high-frequency hearing deficits. The SSN_BB_ and ICRA5-250_BB_ SRT test may thus be complementary in identifying low and high-frequency hearing deficits, respectively.

#### 3.9.2. Origin of Speech Recognition Deficits

LP and HP portions of speech in SSN or ICRA5-250 noise were not equally affected by ageing and hearing sensitivity. On the basis of the performed SII predictions, observed group differences and individual differences within the older listener group, Fig.9 summarizes the identified mechanisms for BB, LP and HP speech recognition. The top line (blue arrows) indicates significant differences in age, PTA or both between the considered groups in the BB, LP or HP analysis. The red arrows indicate significant SRT or MR changes across the groups, while green bars reflected no significant differences between the group results. The significance p was set to p<=0.016 to include a Bonferroni correction for the number of groups.

**Figure 8.**
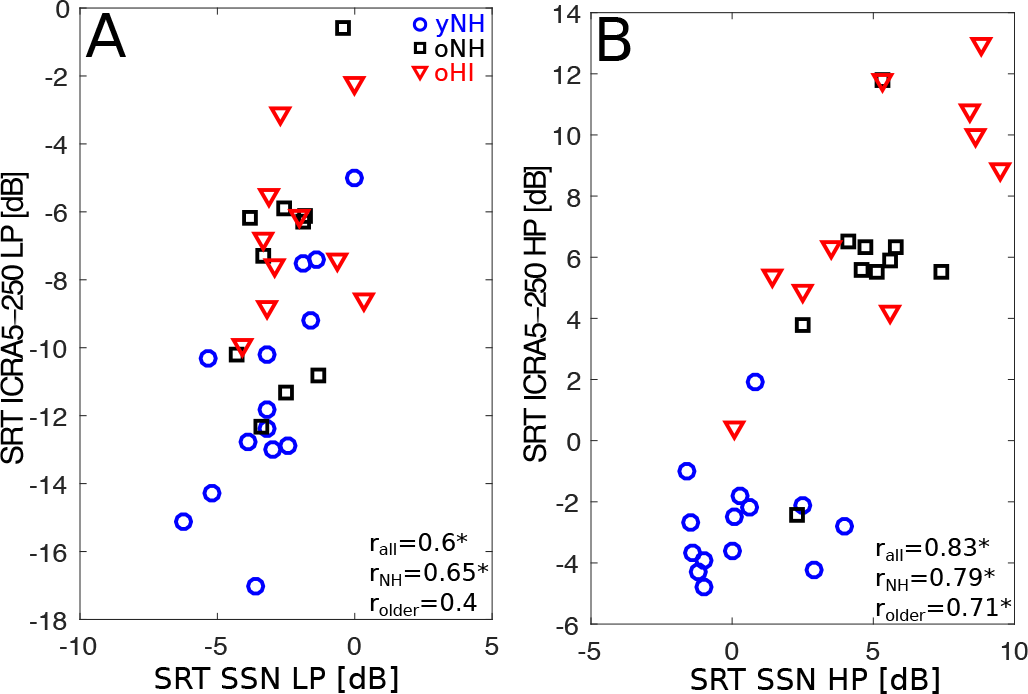
Relations between SRTs in SSN and ICRA5-250 noise. Pearsons’ *r* is given for all participants together (*r*_all_), and for subgroups of normal-hearing (*r*_NH_; yNH+oNH), and older listeners (*r*_older_; oNH+oHI). Significance is indicated by a * for the Bonferroni corrected of p=0.005.

**Figure 9.**
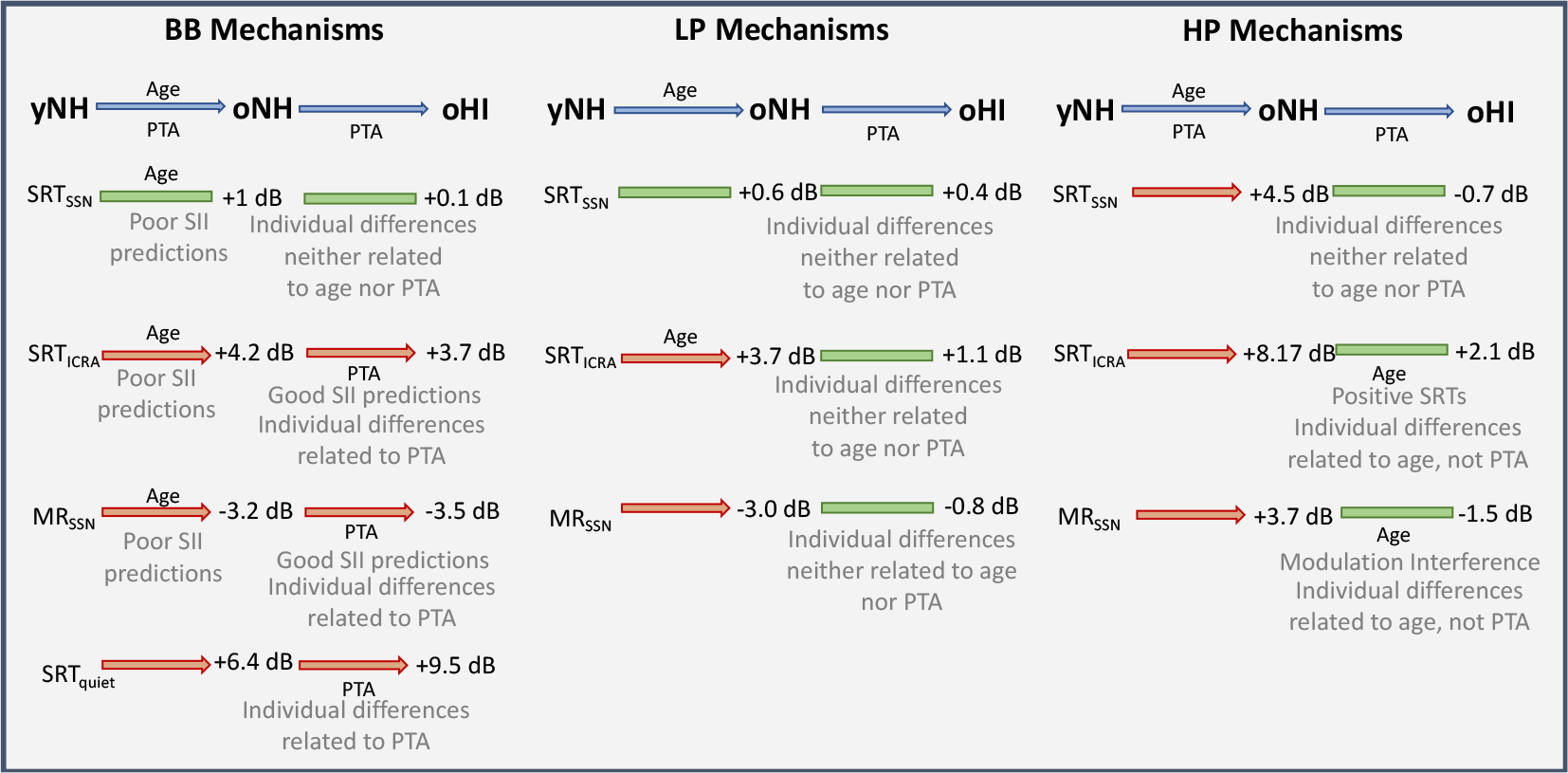
Identified mechanisms degrading SRT and MR performance in yNH, oNH and oHI listeners. Mechanism were identified by combining SII predictions with observed group differences and individual differences within the older (oNH+oHI) listener group for which both age and PTA were normally distributed. The blue arrows indicate which of two factors (age or PTA) were significantly different between the groups. Red arrows indicate significant changes (p<0.016) in group performance, green arrows indicate non-significant changes.

Even though oNH listeners significantly differed in PTA_0.5,1,2,4_ and age from yNH listeners, the SII did not predict their SSN_BB_, ICRA5-250_BB_ SRTs or MR_BB_ performance. This observation corroborates findings from multiple other studies which reported degraded speech recognition in stationary noise (Dubno, Dirks, and Morgan 1984; Füllgrabe, Moore, and Stone 2015; Goossens et al. 2017) and reduced masking release (Dubno, Horwitz, and Ahlstrom 2002; Grose, Mamo, and Hall III 2009; Goossens et al. 2017) for older listeners with normal audiograms. We conclude that there is a mechanism related to increased age which reduces BB speech recognition in oNH listeners. Given that we grouped our listeners into a younger and older group, we were not able to study whether individual differences in performance in the NH subgroup were linearly related to age or were indicative of other (age-related) suprathreshold hearing deficits. When hearing sensitivity worsened in aged listeners, PTA differences became responsible for explaining individual differences in ICRA5-250_BB_ and MR_BB_, while neither PTA (nor age differences) were responsible for explaining individual variations in SSN_BB_. Another supra-treshold hearing deficit must hence be responsible for the reported individual variability (e.g., TFS, TENV or synaptopathy deficits). The origin of these individual differences might be mostly associated with LP mechanisms as SSN_LP_ performance was shown to predict the individual SSN_BB_ performance well (Fig.5), and was shown to be 3 dB better than the SSN_HP_ performance associated with high-frequency coding mechanisms. Even though other studies have identified a low-frequency TFS coding mechanism which can be degraded independently of hearing sensitivity (Füllgrabe, Berthommier, and Lorenzi 2006; Strelcyk and Dau 2009; Grose and Mamo 2010; Hopkins and Moore 2011; Papakonstantinou, Strelcyk, and Dau 2011), we did not assess TFS sensitivity further in the tested listeners so we cannot provide additional support for the origin of the SSN_LP_ variability.

The LP and HP conditions indicated that an overall age-related deficit was responsible for degrading ICRA5-250_BB_ and ICRA5-250_LP_ SRTs in oNH listeners. It is not clear from our analysis whether the oNH ICRA5-250_HP_ SRTs were degraded because of age or PTA differences, but within the older listener group, individual differences were related to age (not PTA) differences.

Lastly, there was a strong relationship between the ICRA5-250_HP_ and SSN_HP_ SRTs within the older listener group, that was absent for the LP conditions (Fig.8). This suggests that within the older group, there is a single HP mechanism degrading performance for both noise types. This mechanism might relate to degraded TENV encoding, which recent studies have associated with age-related synaptopathy in rodents (Parthasarathy and Kujawa 2018). In contrast, we observed that LP speech cues were degraded differently depending on the noise type in the same older listener group.

### 3.10. Masking Release Mechanisms

Negative MR, or modulation interference, has earlier been reported for NH listeners in consonant recognition (Kwon and Turner 2001), for NH listeners performing at positive SRTs (Oxenham and Simonson 2009), and for HI listeners who had positive SRTs (Christiansen and Dau 2012; Goossens et al. 2017). MR_HP_ amounts were shown to be age-dependent in the older listener and given that MR was additive across frequency (i.e., MR_BB_ = MR_LP_ + MR_HP_), there is a benefit in avoiding modulated background noise for the HF regions where older listeners require positive SNRs to reach the SRT.

Lastly, we found that MR was not related to the SSN performance itself, as postulated in other studies (Bernstein and Grant 2009; Christiansen and Dau 2012; Léger, Moore, and Lorenzi 2012b). Instead, we found that MR was significantly related to ICRA5-250 in all LP, HP and BB conditions across either all, older or normal-hearing participant groups with a minimum r of −0.73 (p¡=0.005). Because the added dips in the noise compromised performance (as a function of age) particularly in those listeners who already performed badly in the SSN condition, we postulate that impaired speech-in-noise encoding on the basis of a single HF mechanism (e.g., TENV_HF_ encoding) might be even more challenged when dips are added to the background noise. Recent studies have shown that speech recognition is not only determined by the amount of audible speech information (i.e., above the individual pure-tone threshold or noise level) but that it is also influenced by the temporal-envelope fluctuations inherent to the noise (Stone et al. 2011; Stone, Füllgrabe, and Moore 2012). According to this concept, the modulations contained in the noise signal may also play an important role when apparent stationary maskers are used. Broader cochlear filtering in HI listeners can result in a reduction of inherent low-frequency noise fluctuations which could explain why older listeners showed no MR. However, since broader cochlear filters are not expected in oNH listeners, this concept alone cannot explain the MR loss observed in that group.

#### 3.10.1. SII predictions

In the BB condition, SII failed to predict impaired performance of oNH listeners. This clearly indicates that the drop of performance of oNH listeners can not be explained by their individual pure tone threshold. Hence, supra-threshold deficits need to be explored and included in the SII model. Another modelling approach which, aside from including elevated pure tone threshold, accounts for supra-threshold deficits associated with reduced intensity discrimination or frequency selectivity was proposed by Schädler et al. (2015; 2016). This type of model, or more biophysical models accounting for auditory-nerve deficits (e.g., Zilany, Bruce, and Carney 2014; Verhulst, Altoè, and Vasilkov 2018), can be used to examine the influence of elevated pure tone thresholds as well as supra-threshold deficits on speech intelligibility in stationary and modulated noise in future studies.

The SII was not designed to predict the influence of LP or HP signal filtering on the SRT. Both tested SII approaches failed to predict the measured data. The original version of the model underestimated the influence of reducing the information in the speech signal to LP or HP frequency regions. For the LP condition, the predicted SRTs were almost equal to the BB condition. For the HP condition, an increase of 0.7 and 2.3 dB was predicted in SSN and ICRA5-250 noise, respectively. These values were considerably smaller than the measured difference between BB and HP conditions (7.2 dB for SSN and 16.1 dB for ICRA5-250). The underestimation of the model may be related to the fact that the remaining information after filtering in the frequency band close to the cut-off frequency leads to high SNR in this band and contributes considerably to speech intelligibility, hence biasing the SII predictions. For this reason, we also tested another approach which used a reduced number of SII frequency bands in the SRT calculation, i.e., including only frequency bands corresponding to the frequency bands of the filtered stimuli. This approach resulted in different trends in the predicted SRTs than for the unmodified SII. Generally, an increase in SRT for both filtered conditions and maskers was observed, but in contrast to the empirical data, the predicted SRTs in the LP condition were higher than in the HP condition. This indicates an overestimation of high-frequency speech information in the model. Similar trends were observed for SRT predictions in listeners with downward-sloping high-frequency hearing loss; elevated threshold at high frequencies resulted in much higher predicted SRTs than observed in the empirical data (Hülsmeier et al. 2018). We conclude that the SII model in its present form does not adequately capture how LP and HP speech regions contribute to the SRT.

## 4. Acknowledgements

The work was supported by the DFG Cluster of Excellence EXC 1077/1 ‘Hearing4all’ (AW/SV) and by the DFG research project No 325439187 (Multilingual model-based rehabilitative audiology). It also received funding from the European Research Council (ERC) under the European Union’s Horizon 2020 research and innovation programme under grant agreement No 678120 RobSpear (SV). We thank Sabrina Pieper for the recruitment of study participants and data collection.

